# DNA methylation landscapes in human cells and their chromatin determinants

**DOI:** 10.1101/2025.11.17.688913

**Authors:** Wei Cui, Zhijun Huang, Gerd P. Pfeifer

## Abstract

DNA methylation patterns are established during development and are propagated in a cell type specific manner, but these patterns may become aberrant during aging and cancer. Regions of alternating high and moderate to low levels of DNA methylation exist along all chromosomes in human cells. It is unclear how these distinct DNA methylation blocks are established. Here we have profiled DNA methylation at single base resolution and various histone modifications in human bronchial epithelial cells. We found that many regions of lower DNA methylation are characterized by presence of the Polycomb repressive complex 2 (PRC2) mark histone H3 K27 trimethylation but less so by the PRC1 mark histone H2A K119 monoubiquitylation. These same PRC2-marked regions also showed a depletion of histone H3K36 di- and tri-methylation. Since H3K36me2 and H3K36me3 are recognized by the reader domains of the DNA methyltransferases DNMT3A and DNMT3B and H3K36 methylation is a block to the PRC2 methyltransferases, these crosstalks explain the stable maintenance and antagonism of H3K27me3 and DNA methylation domains. The data give insight into how DNA methylation patterns are established in human cells. We discuss these findings and their potential relevance for altered DNA methylation patterns seen in aging tissues and in cancer cells.

## INTRODUCTION

Somatic cells and tissues of mammalian organisms, with few exceptions, contain the same genome in terms of DNA sequence. Yet, there are hundreds of phenotypically different cell types, which raises the question how such diversity can arise independent of the primary DNA sequence. In each cell, DNA is packaged into chromatin, a structure that consists of DNA wrapped around nucleosome core particles which are further modified by a series of posttranslational histone modifications. The DNA itself also is modified and carries methyl groups or hydroxymethyl groups, chiefly at CpG (CG) DNA sequences where the position 5 of the cytosine ring is modified. Collectively, the assembly of modified DNA and histones, along with the three-dimensional packaging of chromosomes into defined territories, compartments and topologically associated domains (TADs) is referred to as the epigenome. The epigenome is inherently plastic, allowing for changes during development and differentiation. Cell type identity is determined by different sets of epigenomes between each cell type. The most important readout of a specific epigenome is the establishment of cell type specific gene expression patterns.

Active genes generally carry a distinct set of defining epigenomic marks. They include an unmethylated, often GC-rich promoter, referred to as a CpG island [1]. At the chromatin level, active promoters are characterized by histone H3 lysine 4 (H3K4) methylation with H3K4 trimethylation (H3K4me3) being a typical promoter mark [2], and histone acetylation, for example H3K27 acetylation. Downstream of transcription start sites, along the gene body, we find other specific marks, DNA CpG hypermethylation, H3K36 trimethylation (H3K36me3) and H3K79 trimethylation [3,4].

Genomic regions that lack active genes are often associated with repressive histone marks. They have much lower histone acetylation and carry the mark H3K9 trimethylation (H3K9me3) [5]. Such regions are referred to as constitutive heterochromatin and are often found in nuclear lamina associated domains (LADs) [6,7]. Some genes are repressed by a more flexible system referred to as facultative heterochromatin [8]. It means that these regions may be active during specific phases of development or in specific cell types but are otherwise repressed. A typical hallmark of facultative heterochromatin is the Polycomb system. It consists of two sub pathways, Polycomb repression complex 1 (PRC1) and Polycomb repression complex 2 (PRC2) [9,10]. These multi-protein complexes contain enzymatic activities that install histone modifications that interfere with gene activity. The most widely studied such activity is the EZH2 (and EZH1) histone methyltransferase activity of the PRC2 complex, which methylates lysine 27 at the N-terminal tail of histone H3 to produce H3K27me3 [9]. The major enzymatic activity of the PRC1complex is the transfer of one ubiquitin moiety onto lysine 119 of histone H2A to produce H2AK119ub1 [11–13]. PRC1 and PRC2 complexes interact structurally and functionally [9]. They promote the compaction of chromatin to make it less accessible to transcription factors and RNA polymerase enzymes.

With the common use of genomic mapping techniques beginning about two decades ago [14–16], much knowledge has been assembled about the specific features of histone modifications and DNA methylation patterns in many different cell types. Through numerous uncoordinated and coordinated (e.g., by ENCODE) efforts, the epigenome of commonly used cell types and mammalian tissues have been mapped comprehensively. Of great interest has always been the question if the epigenome is aberrant during the initiation of human diseases or during the aging process. For example, how would an altered epigenome that is found in one cell type promote a tissue-specific disease and why would this alteration be tissue-specific? Through the work of countless laboratories, we now know that cancer cells are characterized by wide-spread epigenomic alterations. This field was initiated over four decades ago when it was reported that cancer cells from animals or humans have altered DNA methylation patterns, either genome-wide [17,18] or at specific gene loci [19]. The methylation changes include genome-wide DNA hypomethylation, which affects many parts of the genome in different cancer types [20], as well as more localized DNA hypermethylation events, which targets mostly CpG islands [21,22]. In some cases, methylation of promoter-based CpG islands may silence critical genes, for instance tumor suppressor genes or genes essential for maintaining the differentiated state of a somatic cell [23–26]. Although studied less widely and less systematically, alterations in chromatin structure are also found in tumors. Most notably, mutations in many cancer types are observed in genes that encode enzymes of the epigenetic machinery, for example histone methyltransferases and acetyltransferases, components of Polycomb complexes and chromatin remodeling factors [27,28].

In addition to cancer and a few other diseases, the aging process also leads to a disruption of epigenomic patterns. This has most widely (in fact almost exclusively) been studied for DNA methylation. Similar to cancer, aging leads to a general loss of DNA methylation and at the same time to an encroachment of DNA methylation into CpG islands [29–31]. Often similar groups of genes are affected in cancer and aging. Most notably, CpG islands targeted by the Polycomb complexes, such as developmental transcription factor genes, are methylated during carcinogenic processes [16,32–34], but also during the aging process [35,36] and during inflammation [37], a condition that may predispose to caner and aging. Since the discovery of an active DNA demethylation process about 15 years ago [38], it has now become clear that the establishment, maintenance and erosion of DNA methylation patterns is much more complicated than previously thought. DNA methylation is established by DNA methyltransferases that methylate CpG sequences [39]. The active enzymes include the maintenance methyltransferase DNMT1 and the de novo methyltransferases DNMT3A and DNMT3B which methylate previously unmethylated sites preferentially. The DNA demethylation process is initiated by oxidation of the methyl group of 5-methylcytosine by the Ten-Eleven-Translocation (TET) enzymes, TET1, TET2, and TET3 [38]. The demethylation process is often defective in human cancer, either by mutation of TET2, or by defects in TET activity [40].

However, what remains largely unclear to this date is how these epigenomic patterns are established mechanistically in normal cells during development and how these patterns become aberrant in cancer. The extent of crosstalk between different epigenomic states and modification patterns along the genome is only partially understood. In our previous work, we reported that dysfunction of the Polycomb protein RYBP, which is a critical component of PRC1 complexes, along with loss of TET proteins leads to widespread hypermethylation of thousands of CpG islands in human bronchial epithelial cells [41]. In that publication, we focused on CpG islands. From the published work and from additional chromatin modification mapping experiments, we have now obtained a detailed picture of how the different epigenomic parameters of DNA and histone modification overlap or antagonize each other. Here, we provide a genome-wide assessment of this crosstalk and show that a tripartite relationship between DNA CpG methylation, H3K27me3 and H3K36 methylation largely shapes long-range epigenomic states along the human genome.

## MATERIALS AND METHODS

### Cell culture

Human bronchial epithelial cells (HBEC-3KT, CRL-4051; RRID: CVCL_X491) [42] were obtained from ATCC. Cell lines were authenticated by ATCC by short tandem repeat profiling. The cells were cultured in keratinocyte serum-free medium (K-SFM; Thermo, 17005042) containing 50 µg/ml bovine pituitary extract and 5 ng/ml human recombinant epidermal growth factor and incubated at 37 °C with 5% CO_2_. The media were changed every 3 to 4 days.

### DNA methylation analysis

Genome-wide DNA methylation patterns at single base resolution were determined by whole genome bisulfite sequencing (WGBS). The analysis pipeline for WGBS and the data obtained have been published previously [41,43,44].

### ChIP-seq

ChIP-seq was performed as previously described [41]. H2AK119ub1 (CST, 8240), H3K27me3 (CST, 9733), H3K36me2 (CST, 2901S), and H3K36me3 (Abcam, ab9050) antibodies were used for immunoprecipitation. Briefly, cells were cross-linked with 1% formaldehyde for 10 min at room temperature, and the reaction was quenched with glycine. Crosslinked cells were lysed in lysis buffer and incubated on ice for another 10 min. The washed cell pellets were resuspended in shearing buffer and sonicated to shear the DNA into 300 to 500 bp long fragments. An optimal amount of chromatin, antibodies, and protein-G bead complexes were incubated overnight on a rotator at 4°C. Purified DNA was quantified for library preparation with the Qubit sensitivity dsDNA HS Kit. Libraries were prepared using the TruSeq ChIP Sample Preparation Kit (Illumina, IP202–1012, IP-202–1024) according to the manufacturer’s instructions. The libraries were pooled and sequenced with an Illumina NextSeq500 or NovaSeq6000 instrument. A 5% spike-in of PhiX DNA was added to the runs to increase diversity and for quality control purposes. Each library had at least 50 million reads. Three replicates were generated in this study.

### ChIP-seq data analysis

ChIP-seq data analysis was performed as previously described [41,43]. Briefly, the adapter and low-quality sequences were trimmed with Trimgalore (v 0.4.0) with default parameters, then the trimmed reads were mapped to the human genome hg19 with Bowtie2 (v 2.3.5). The deduplicated reads were used for peak calling by HOMER (v 4.10). Density plots of ChIP-seq enrichment at peak center regions were made with the R package genomation (version 1.28.0). For all overlap analysis of ChIP peaks, we used the bedtools intersection function, as well as for all other region overlap analysis in this study.

## RESULTS

### DNA methylation is segregated into highly methylated and partially methylated regions along the genome

DNA methylation was analyzed at single base resolution by the WGBS method as described previously [41,43]. Triplicate samples of bronchial epithelial cell clones of the telomerase- and CDK4-immortalized cell line HBEC3 were used. Broader genomic regions at the scale of one to several megabases were inspected as shown in Figure 1 for several chromosomal segments. Most genomic regions are subdivided into segments of high DNA methylation (over 80% at most CpG sites) and intermediate or partial methylation (30-60%). Using genome-wide analysis of genic and intergenic regions, we observed the highest DNA methylation levels in genic regions, also referred to as gene bodies (Fig. 2A). This is consistent with prior work in other cell types [45–47]. Methylation levels are known to correlate with gene expression levels [45,48].

**Fig. 1.**
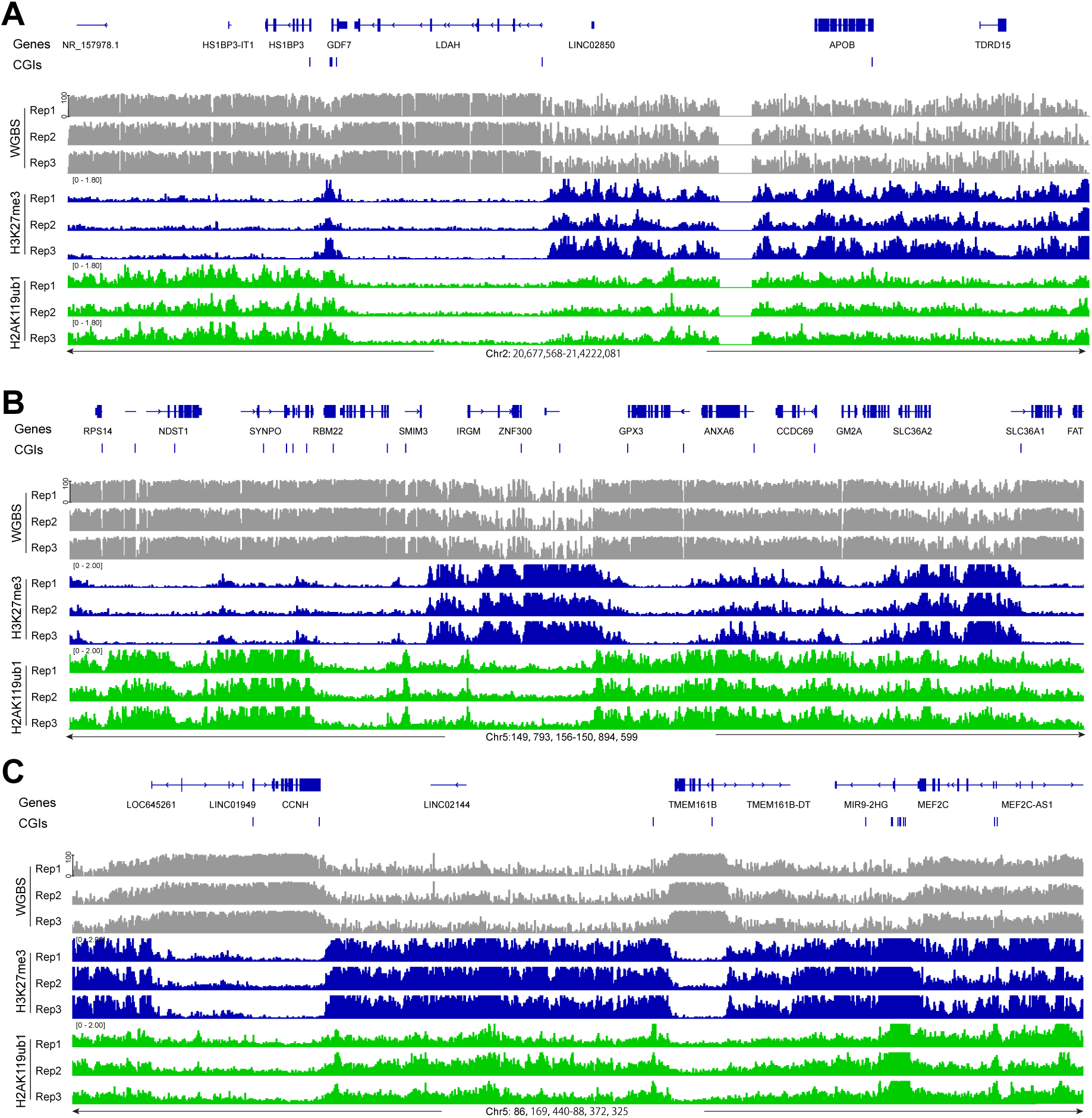
Display of DNA methylation at PRC1- and PRC2-marked regions. **A.** DNA methylation (grey), H3K27me3 ChIP-seq (blue), and H2AK119ub1 (green) ChIP-seq tracks for a representative region of chromosome 2. Hg19 coordinates are shown along with the CpG island regions (blue bars). **B.** DNA methylation, H3K27me3 ChIP-seq, and H2AK119ub1 ChIP-seq tracks for a representative region of chromosome 5 (positions 149,793,156 to 150,894,599). **C.** DNA methylation, H3K27me3 ChIP-seq, and H2AK119ub1 ChIP-seq tracks for a second representative region of chromosome 5 (positions 86,169,440 to 88,372,325).

**Fig. 2.**
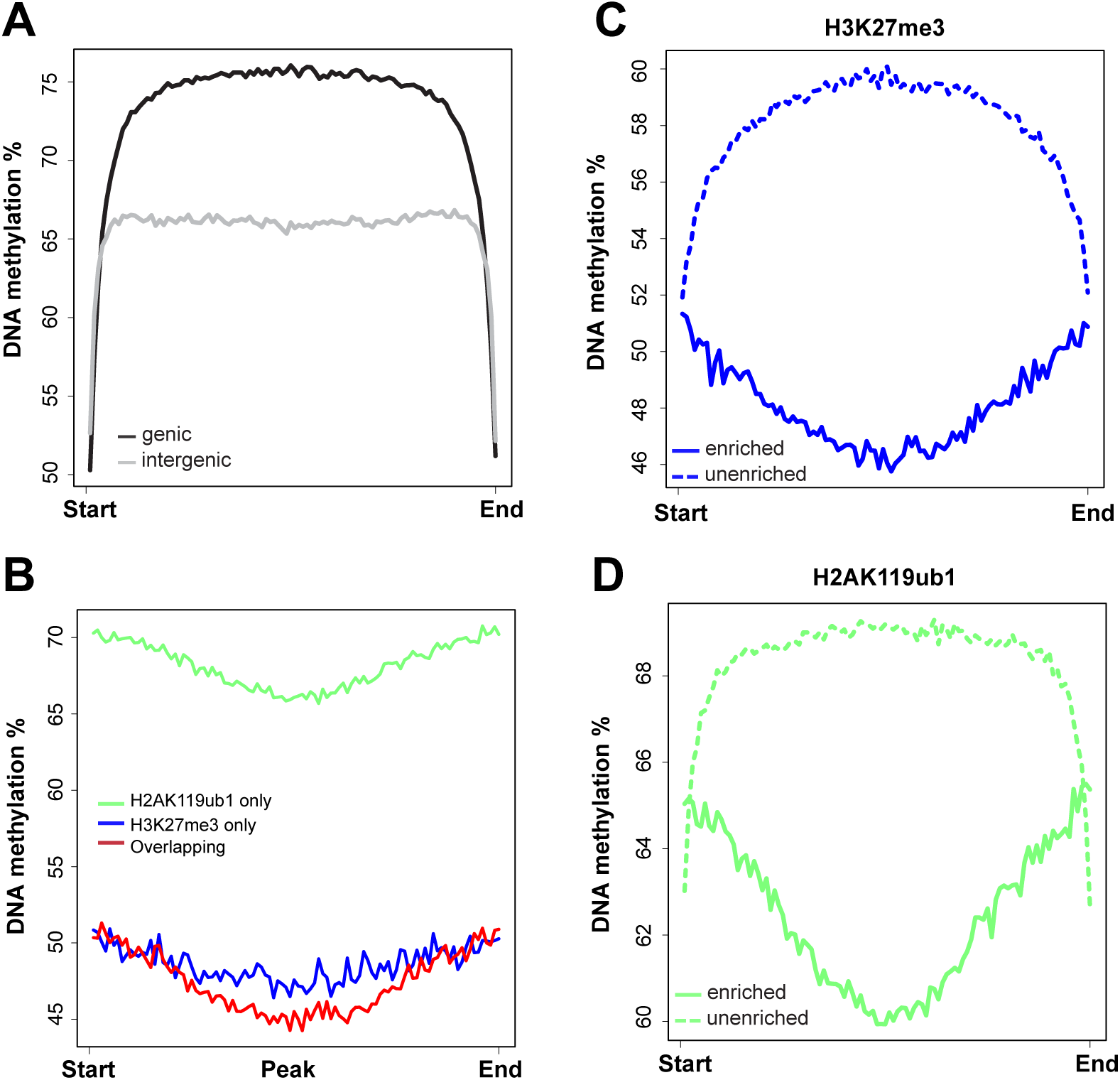
The profile of DNA methylation at PRC1- and PRC2-targeted regions. **A.** Meta-signal plots of DNA methylation at genic and intergenic regions. **B.** Meta-signal plots of DNA methylation at H2AK119ub1-only, H3K27me3-only, and at H2AK119ub1 and H3K27me3 broad peak overlapping regions. **C.** Meta-signal plots of DNA methylation at all H3K27me3-enriched and unenriched regions. **D.** Meta-signal plots of DNA methylation at all H2AK119ub1-enriched and unenriched regions.

We also find that many of the partially methylated domains fall into gene-poor areas of the genome. Genome segments that are depleted of active genes tend to be associated with the nuclear lamina [49]. Although we lack data on lamina-associated domains (LADs) in this bronchial cell line, constitutive LADs are often common between different human cell types. The reason why LADs have lower DNA methylation levels have been debated. Suggestions include late DNA replication of these domains allowing less time for DNA methyltransferases to complete their reactions before mitosis [50], a sequence preference for DNMTs that disfavors A/T-CG-A/T DNA sequences [51], and as we proposed recently, a limited access of DNMT enzymes to regions attached to the nuclear lamina [43]. Despite the clear segregation of the DNA methylation patterns into highly methylated and partially methylated domains, it has remained unknown how these bipartite patterns are established and maintained along the genome.

### Lower DNA methylation is best correlated with the PRC2 mark, H3K27me3

In parallel with DNA methylation, we mapped the distribution of various histone marks in the same cell clones. We focused here on the Polycomb marks H3K27me3 installed by the EZH1/2 histone methyltransferases of the PRC2 complex and on H2AK119ub1 installed by the PRC1 complex. As can be seen in Figure 1 and several subsequent Figures, the presence of strong broad peaks of H3K27me3 generally coincides with regions of lower DNA methylation. When we compared DNA methylation levels over regions covered by H3K27me3 broad peaks, H2AK119ub1 broad peaks or broad peaks containing both marks, we observed that DNA methylation levels are higher at H2AK119ub1 marked regions but much lower at genomic segments marked by H3K27me3 or by both marks (Fig. 1; Fig. 2B-D).

### H3K27me3 is reverse to H3K36me2 and H3K36me3

Next, we determined the genomic distribution of the H3K36me2 and H3K36me3 histone modifications (Fig. 3). These marks are established by the NSD family of histone methyltransferases, NSD1, NSD2, and NSD3 [52]. H3K36me3 produced by SETD2 is a mark preferentially found in gene bodies where it likely accumulates in conjunction with RNA polymerase II progression [53]. It is thought that H3K36me3 promotes gene activity by suppressing intragenic initiation and antisense transcription. Genomic profile plots in Figure 3A show that H3K36me3 is indeed accumulating in gene bodies genome-wide as opposed to intergenic regions. In contrast, H3K36me2 is more broadly distributed and is found along genes and in non-genic regions (Fig. 3A). In Figure 3B, we establish the profiles of the Polycomb marks H3K27me3 and H2AK119ub1 along genic and intergenic regions. Genome-wide, the H2AK119 ubiquitin mark occurs in genes and in non-genic regions but H3K27me3 is more abundant in intergenic segments compared to genes (Fig. 3B). An example of that is shown in Figure 3C. Figure 3C shows a part of chromosome 5 with the distribution of these chromatin marks and that of DNA methylation determined by WGBS. The preferred distribution of H3K36me3 along genes can easily be recognized (Fig. 3A, 3C) whereas H3K36me2 is found at genes and in between genes. This Figure also shows a noticeable reduction of histone H3 K36 modifications in an area with strong marking by the Polycomb system, as shown by the strong enrichment patterns of H3K27me3 and H2AK119ub1 (right side of Figure 3C).

**Fig. 3.**
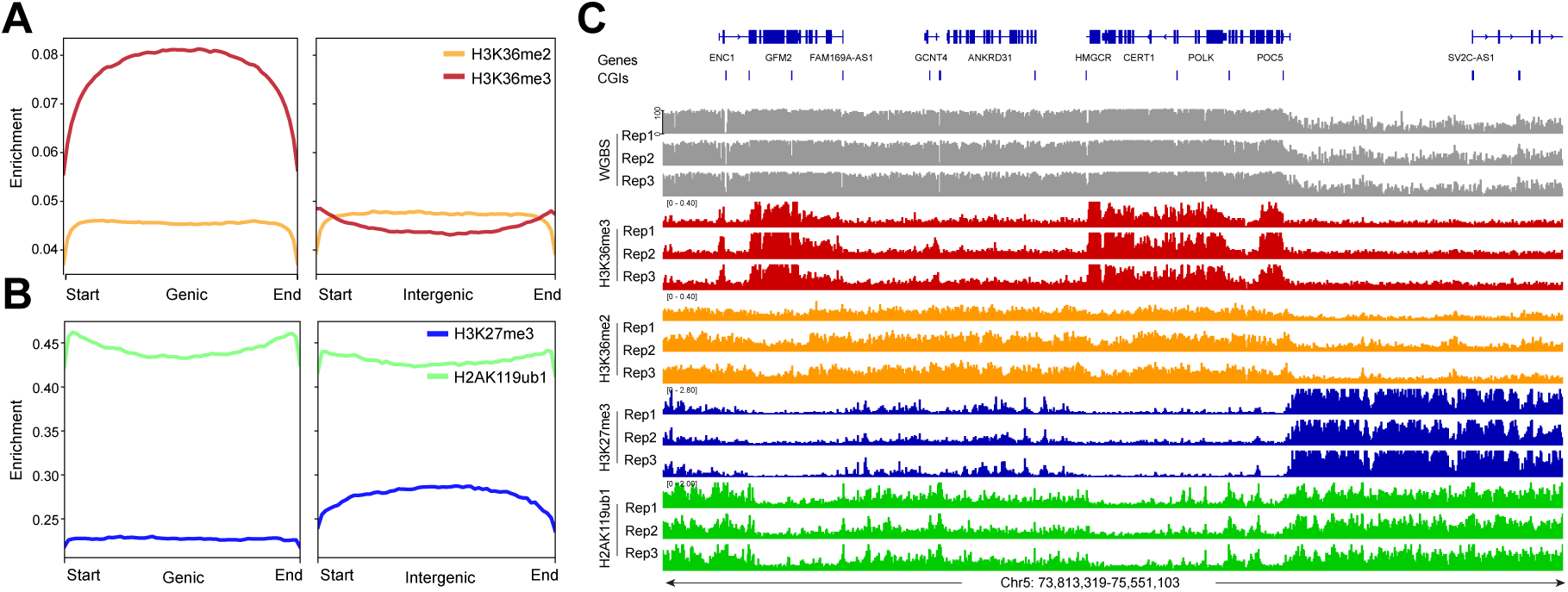
The enrichment of H3K36me2, H3K36me3 and the Polycomb marks at genic and intergenic regions. **A.** Meta-signal plots of H3K36me2 and H3K36me3 enrichment at genic and intergenic regions in HBEC3 cells. **B.** Meta-signal plots of H3K27me3 and H2AK119ub1 enrichment at genic and intergenic regions in HBEC3 cells. **C.** DNA methylation (grey), H3K36me3 ChIP-seq (red), H3K36me2 ChIP-seq (orange), H3K27me3 ChIP-seq (blue), H2AK119ub1 ChIP-seq (green) tracks for a representative region at chromosome 5.

We next investigated the crosstalk between the Polycomb marks and the H3K36 methylation system (Fig. 4; Fig. 5). It is easily recognizable that regions with high levels of H3K36me3 have extremely low levels of H3K27me3 or H2AK119ub1 (Fig. 4A). When there are broad peaks of H3K36me2, the Polycomb marks are also reduced. Vice versa, genomic areas that are strongly marked by H3K27me3 and H2AK119ub1 show low levels of the H3K36 modifications. Figures 4B and 4C show a correlation profile analysis. At the genome-wide levels, regions bound by the Polycomb marks show lower levels of H3K36me3 (Fig. 4B) and H3K36me2 (Fig. 4C). Figure 5 shows additional examples of this relationship. Profile plots of H2AK119ub1 broad peaks show some enrichment for H3K36me3 (Fig. 5A) and H3K36me2 (Fig. 5B). However, when H3K27me3 is present alone or when it is overlapping with H2AK119ub1, the enrichment of the H3K36 modifications is much less.

**Fig. 4.**
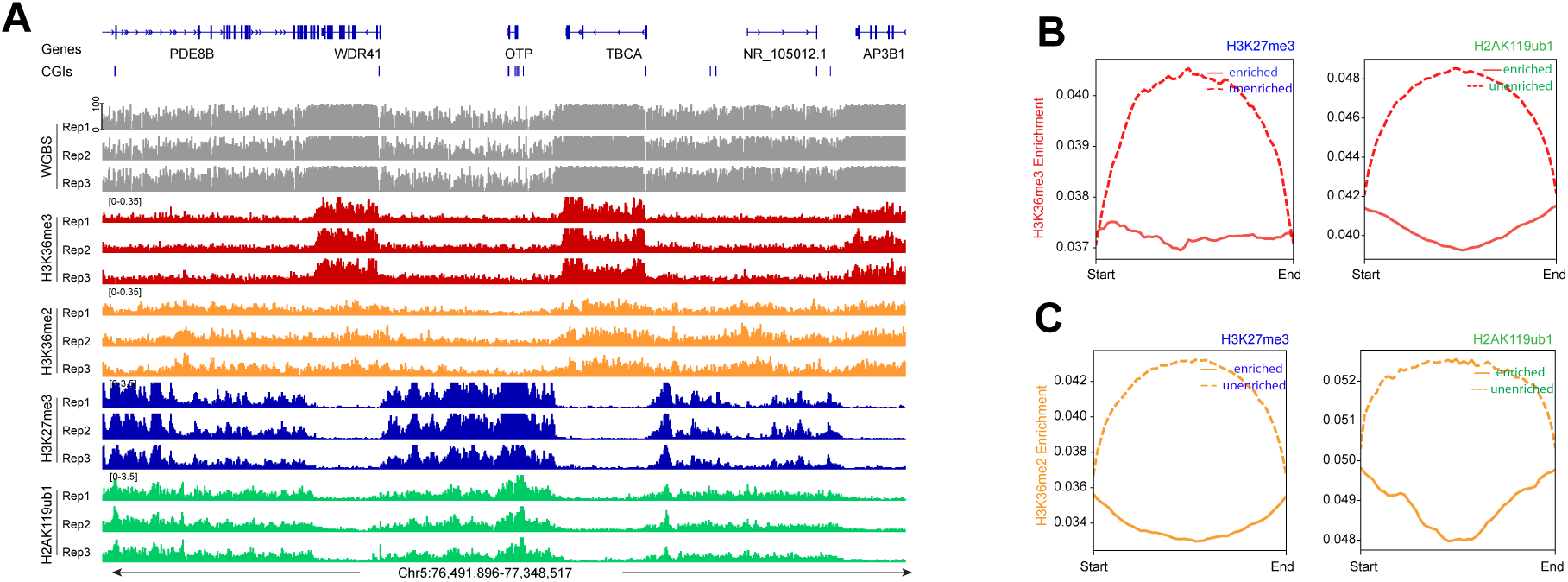
H3K36me2, H3K36me3 and Polycomb marks show reverse occupancy. **A.** DNA methylation, H3K36me3 ChIP-seq, H3K36me2 ChIP-seq, H3K27me3 ChIP-seq, and H2AK119ub1 ChIP-seq tracks for a representative region at chromosome 5. **B.** Meta-signal plots showing ChIP-seq signal for H3K36me3 at PRC-targeted and -untargeted regions in HBEC3 cells, respectively. **C.** Meta-signal plots showing ChIP-seq signal for H3K36me2 at PRC-targeted and -untargeted regions in HBEC3 cells, respectively.

**Fig. 5.**
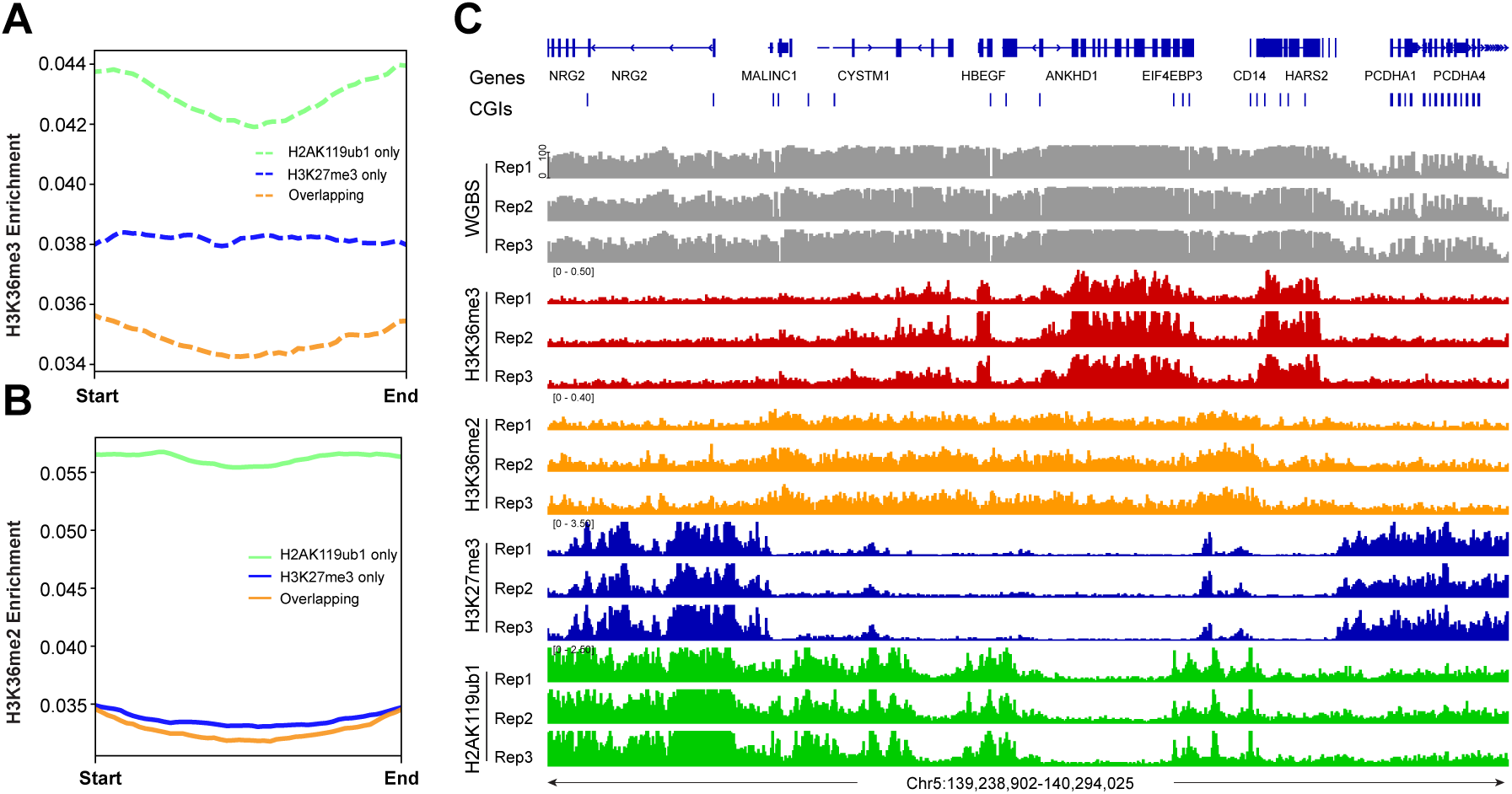
The profile of H3K36me2 and H3K36me3 at all PRC1- and PRC2-marked regions. **A.** Meta-signal plots showing ChIP-seq signals for H3K36me3 at H2AK119ub1-only, H3K27me3-only, and their overlapping regions in HBEC3 cells. **B.** Meta-signal plots showing ChIP-seq signals for H3K36me2 at H2AK119ub1-only, H3K27me3-only, and their overlapping regions in HBEC3 cells. **C.** DNA methylation, H3K36me3 ChIP-seq, H3K36me2 ChIP-seq, H3K27me3 ChIP-seq, and H2AK119ub1 ChIP-seq for a representative region at chromosome 5.

### High DNA methylation levels are linked to H3K36me2 genome-wide

When we integrated the distribution of DNA methylation versus the different histone modification profiles, we found that DNA methylation best correlates with H3K36me3 levels, and then also correlates with H3K36me2, the latter not being limited to gene bodies but more broadly distributed (Fig. 6; Fig. 3C; Fig. 5C).

**Fig. 6.**
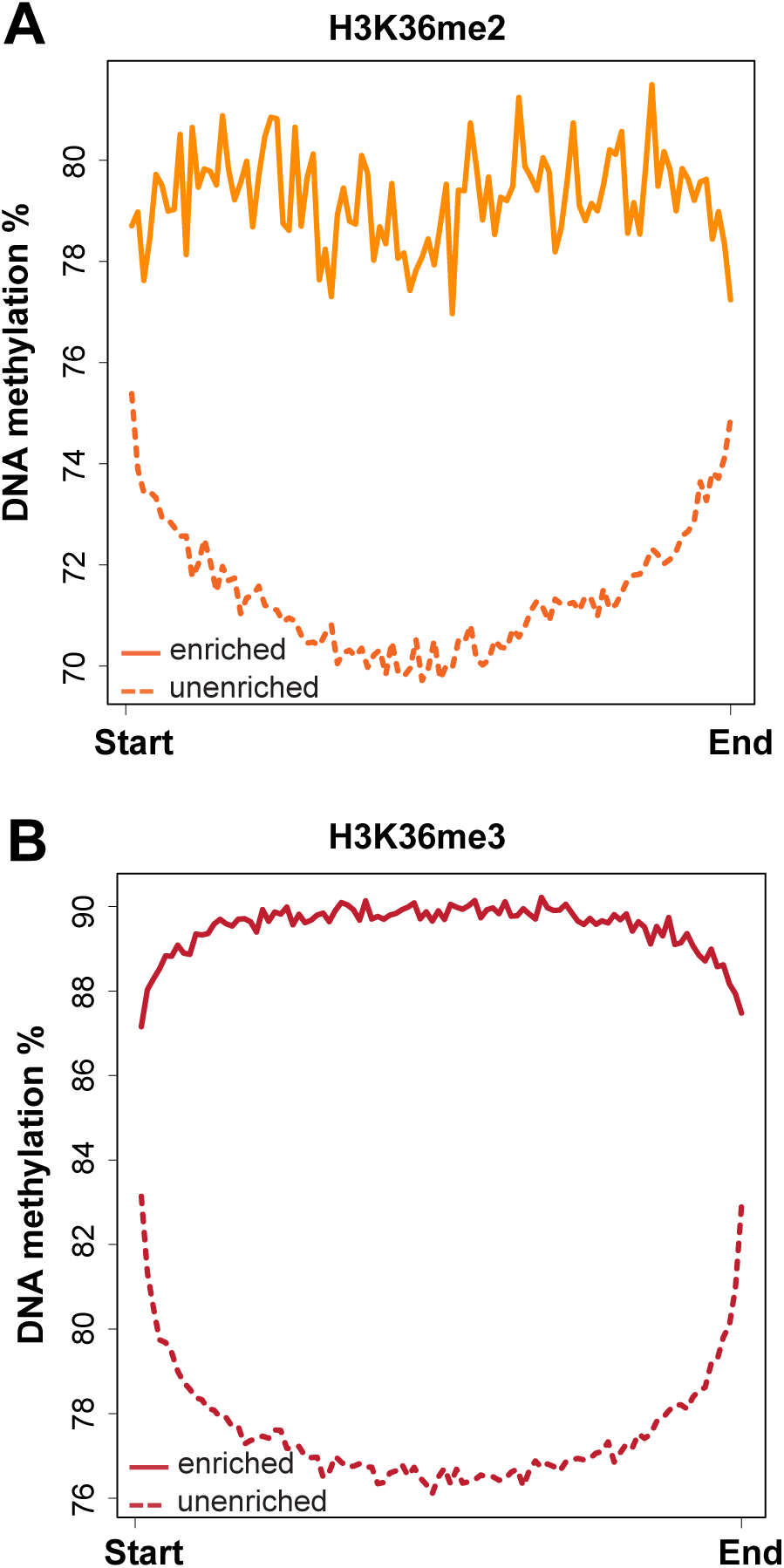
DNA methylation at H3K36me2- and H3K36me3-marked regions. **A.** DNA methylation is higher at genomic regions enriched for H3K36me2. **B.** DNA methylation is highest at genomic regions enriched for H3K36me3.

### DNA methylation at CpG islands

In the last part of our analysis, we focused on CpG islands, CpG-rich genomic regions, commonly found at promoters that are often unmethylated when the gene is active. Indeed, most CpG islands of the genome have relatively low DNA methylation levels (Fig. 7A) even though some of these regions are at least partially methylated, for example when they occur in gene bodies or in intergenic regions. We then profiled the histone marks over all CpG islands. Both Polycomb marks can occur at CpG islands, with higher enrichment for H2AK119ub1 (Fig. 7B). Interestingly, these Polycomb marks show higher levels at CGI borders. In contrast, H3K36me2/3 are quite strongly depleted at CpG islands with the lowest levels found for H3K36me2 (Fig. 7C). To score differences between CpG islands that have low or high DNA methylation, we separated the CGIs into groups containing either CGIs with less than 25% methylation or CGIs with more than 75% methylation (Fig. 7B, 7C) and then profiled the histone marks. This analysis showed again very low levels of H3K36me2 in both categories but higher levels of H3K36me3 at the more highly methylated CGIs. Most of these H3K36-me3-marked CGIs were localized in transcribed gene bodies. These regions are preferentially targeted by de novo DNA methylation activities [54]. We show two examples; one is for a methylated CpG island at the *NKX2-4* gene (Fig. 7D), which is not transcribed in HBEC3 cells, and one for an unmethylated CpG island adjacent to the gene *ATG12* (Fig. 7E). Although there is some H2AK119ub1 modification over these regions, these CpG islands are generally well depleted of H3K36me2 and H3K36me3. This data suggest that methylated CpG islands may show a different relationship between DNA methylation and H3K36me2 deposition compared to non-CpG island regions.

**Fig. 7.**
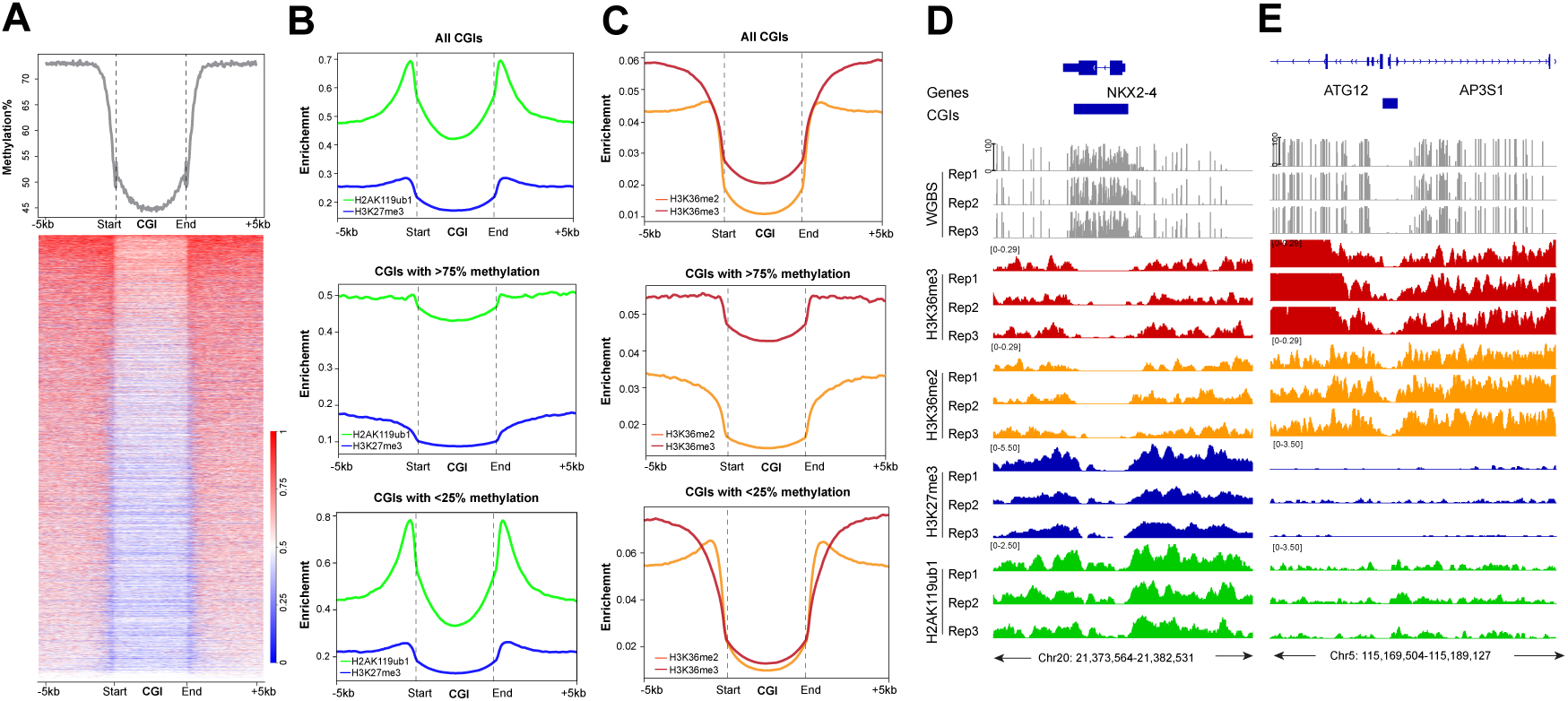
The profile of DNA methylation and histone marks at CpG islands. **A.** Meta-signal plot and heatmap of DNA methylation at all CpG islands (CGIs) and 5-kb flanking regions in HBEC3 cells. The heatmap shows DNA methylation levels of each CGI and 5-kb flanking regions. Three biological replicates were analyzed. **B.** Meta-signal plots of H2AK119ub1 (green) and H3K27me3 (blue) ChIP-seq signals at all CGIs, CGIs with >75% methylation, and CGIs with <25% methylation levels, respectively, including 5-kb flanking regions. **C.** Meta-signal plots of H3K36me2 (orange) and H3K36me3 (red) ChIP-seq signals at all genomic CGIs, CGIs with >75% methylation, or <25% methylation levels, respectively, including 5-kb flanking regions. **D.** DNA methylation (gray), H3K36me3 ChIP-seq (red), H3K36me2 ChIP-seq (orange), H3K27me3 ChIP-seq (blue), and H2AK119ub1 ChIP-seq (green) in HBEC3 cells at the CpG-methylated *NKX2-4* gene locus. Hg19 coordinates are shown along with the CGI region (blue bar). **E.** DNA methylation (gray), H3K36me3 ChIP-seq (red), H3K36me2 ChIP-seq (orange), H3K27me3 ChIP-seq (blue), and H2AK119ub1 ChIP-seq (green) in HBEC3 cells at the unmethylated *ATG12* gene locus. Hg19 coordinates are shown along with the CGI region (blue bar).

## DISCUSSION

In this study, we established links between genome-wide DNA methylation profiles and four prominent histone modifications in a human bronchial epithelial cell line. We did not focus here on the well-known relationship between the promoter mark H3K4me3, which interferes with de novo methylation by DNA methyltransferases [55] and is critical for keeping active and poised promoter CpG islands free of DNA methylation. The antagonistic relationship between DNA methylation and the Polycomb mark H3K27me3 has previously been noted [30,56–61]. In some experimental system, for example during the progression of normal cells to malignancy, there can be a switch of the two repressive marks. Loss of the Polycomb mark at target genes may occur in cancer when it is replaced with DNA methylation [37,62]. In the opposite direction, when DNA methylation is reduced, for example artificially by treatment of cells with a DNA methylation inhibitor, the demethylated regions may become occupied and modified by the Polycomb system [63]. In these situations, DNA methylation and Polycomb marking appear as opposite events, similar as we have observed in our experimental system (Fig. 1). The reason why this exclusion occurs is unclear. It is possible that highly CpG-methylated DNA interferes with Polycomb deposition since PRC complexes contain subunits (for example, KDM2A, KDM2B) that prefer binding to unmethylated CpG-rich DNA through their CXXC zinc finger domains [64–67]. On the other hand, once a region is occupied by Polycomb and the histones are marked by these activities, these complexes and modifications may not allow access of DNA methyltransferases.

However, our data suggest that the situation is more complicated because of additional crosstalk between specific histone modifications and between DNMTs and those histone modifications. First, the histone modifications on the histone H3 N-terminal tail are themselves mutually exclusive. The EZH2-catalyzed H3K27 methylation activity of PRC2 is sensitive to the methylation state of the H3K36 lysine residue, at least when present on the same histone tail [68]. The activity of EZH2 is much higher on H3 histone tails that are unmodified at lysine 36 compared to dimethylated or trimethylated H3K36 [69,70]. Cryo electron microscopy studies have shown that histone H3 tails containing H3K36me3 interact poorly with PRC2 [71].

The methylation marks at H3K36 are dimethylation established primarily by NSD1, NSD2 and NSD3 as well as trimethylation installed by the SETD2 methyltransferase [72]. However, H3K27me3 is not known to block NSD or SETD2 activity, although loss of NSD1 leads to genome-wide expansion of H3K27me3 domains [73]. A mutually exclusive domain model may explain the stability of histone H3K27 and H3K36 histone methylation landscapes [74]. These inhibition models suggest that once an active domain with H3K36me2 and even more so with H3K36me3 has been established it is difficult to overcome this active state by PRC-mediated Polycomb repression.

The H3K36 histone modifications explain not only the absence of H3K27 repressive marks at these regions but also the antagonism between H3K27 methylation and DNA methylation. This triangular relationship (Fig. 8) exists because the H3K36 methylation marks positively induce DNA methylation. This was first shown for the de novo DNA methyltransferase DNMT3A which interacts through its PWWP reader domain with methylated H3K36 [75,76]. DNMT3B, which preferentially resides in gene bodies, also binds through its PWWP domain to H3K36-modified histones and prefers H3K36me3 [77–80]. This finding may explain the high DNA methylation levels present in gene bodies where this histone mark is prevalent. Later it was shown that H3K36me2 is preferentially recognized by the PWWP domain of DNMT3A [81]. Since H3K36me2 is broadly present along many parts of the genome, this modification can explain much of the genomic DNA methylation patterns. Knockout of H3K36me2 methylation led to globally reduced levels of DNA methylation [81]. Interestingly, levels of H3K36me2 are also low in LAD regions, which provides yet another explanation why LADs contain partially methylated domains [43]. Switch of LAD-associated genomic B compartments that contain heterochromatin to active euchromatic A compartments leads to an enhancement of H3K36me2 levels in the switched regions and to the acquisition of long stretches of DNA hypermethylation [43].

**Fig. 8.**
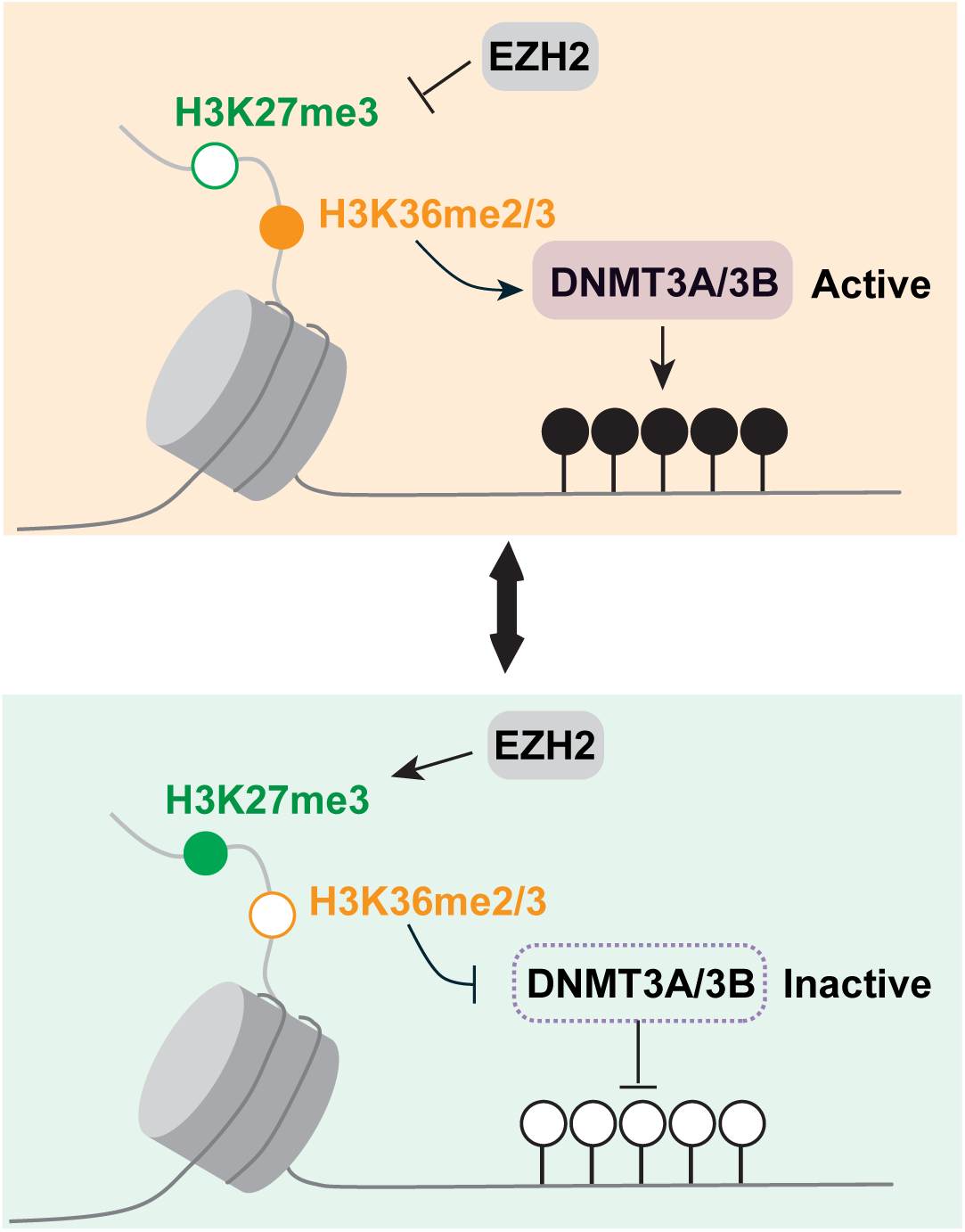
Model of the crosstalk between H3K27me3, H3K36 methylation and DNA methylation. The state of H3K36 methylation determines if H3K27 methylation by EZH2 occurs and if DNA methylation by DNMT3A or DNMT3B takes place.

The question may be asked in which order these genome modifications are established and what it takes to overcome and shift these landscapes. The issues of cause or consequence are often difficult to discern in biological systems with steady state observations. However, the available evidence from in vitro systems and from genetic manipulation experiments would suggest that the NSD and SETD2-catalyzed marks are the primary determinants of both the Polycomb PRC2 mark setting and of the DNA methylation setting (Fig. 8). Therefore, interference with the H3K36 methylation system may be an attractive approach to manipulate either DNA methylation – causing DNA hypomethylation – or to affect the Polycomb system – causing increased Polycomb-mediated repression. However, these two outcomes may neutralize each other in terms of their ultimate effect on gene expression.

One phenomenon that is widely prevalent in most cancer types and is found during the aging process as well is the methylation of Polycomb target genes. In this process, hundreds or even thousands of genes that are marked by the Polycomb system in normal cells, either in embryonic stem cells or in the corresponding normal tissue cells, undergo DNA hypermethylation at CpG islands. Even though this process has been observed almost two decades ago [16,32,33,82,83], the mechanisms leading to this widespread DNA hypermethylation have remained elusive. The same process operates during tissue inflammation [37,84] and during aging [30,35,36,85,86]. Our hypothesis is that Polycomb-marked CpG islands are normally protected from the DNA methylation machinery. One such pathway that protects from DNA methylation Is oxidation of 5-methylcytosine by the 5-methylcytosine oxidases, the TET proteins. TET proteins are often dysfunctional in human cancer due to either mutation or to reduced enzymatic activity [40,87,88]. This phenomenon can easily be observed by the almost complete absence of the primary TET oxidation product 5-hydroxymethylcytosine in human tumors of many different tissue origins [89]. Another protective mechanism against DNA methylation may be the presence of Polycomb complexes themselves at CpG islands. Support for a protection mechanism came from a recent study, in which we inactivated all three TET genes (TET1, TET2, and Tet3) along with the critical Polycomb component RYBP in human bronchial epithelial cells [41]. This manipulation led to DNA hypermethylation at thousands of CpG islands in the gene-modified cells and conferred a tumorigenic phenotype onto the cells. A large fraction of the hypermethylated regions corresponded to the hypermethylated CpG islands found in human squamous cell lung carcinomas [41].

Given the widespread crosstalk between H3K27 methylation, H3K36 methylation and DNA methylation, one wonders if the H3K36 methylation system plays any role in the methylation of Polycomb target genes. CpG islands are bound by the H3K36 demethylase enzymes, KDM2A and KDM2B [64,65], which should reduce these modifications at unmethylated CpG islands consistent with our findings (Fig. 7). The H3K36 modifications are also largely absent from methylated CpG islands (Fig. 7C, 7D) except for H3K36me3 at the bodies of active genes. Is it possible, however, that NSD enzyme activities could forcefully introduce H3K36 methylation at Polycomb-marked CpG islands, which would interfere with PRC2 but promote de novo DNA methylation. In fact, NSD2 and NSD3 are upregulated at the transcript levels or carry activating mutations in human tumors [90–94]. Further work is required to better understand how these mutually dependent genome modification systems are operating in different cell types and how they break down during aging and cancer.

## Author Contribution

Cui, W.: performed experiments and analyzed data.

Huang, Z.: analyzed data.

Pfeifer, G.P.: conceptualized the study, wrote the first draft of the manuscript and acquired funding.

All authors participated in manuscript revisions.

## Ethical Approval

Not applicable.

## Consent to Participate

Not applicable.

## Consent for Publication

Not applicable.

